# Mon1-Rab7 axis is essential for transport, localization and anchoring of *oskar* mRNA

**DOI:** 10.64898/2025.12.24.696314

**Authors:** Vasudha Dwivedi, Vrushali Katagade, Sourav Halder, Jyotish Sudhakaran, T Anjana, Girish S Ratnaparkhi, Vasudevan Seshadri, Anuradha Ratnaparkhi

## Abstract

Asymmetric localization of *oskar* mRNA to the posterior of the oocyte is a complex process driven by autonomous and non-cell autonomous mechanisms. The former includes Oskar protein that reinforces localization and anchoring of its mRNA through activation of endocytosis and regulation of actin cytoskeleton; the latter includes signals from the posterior follicle cells (PFCs) that regulates microtubule orientation for polarized transport. Here we identify Monensin Sensitivity 1 (Mon1), as a novel factor regulating anterior-posterior (A-P) patterning. Mon1 is an evolutionarily conserved activator of Rab7-a key regulator of the endo-lysosomal pathway. Embryos lacking maternal *mon*1 (*mon1^m^*) show mislocalized *oskar* and *bicoid* mRNAs leading to loss of patterning and lethality. In the mutant oocyte Staufen appears clumpy and the levels of Oskar protein and Par-1 is significantly reduced. Abnormal actin rings are seen in the ooplasm. Driving expression of *mon1* in the germline rescues these phenotypes and restores viability. In contrast, expression in the PFC predominantly rescues the Par-1 phenotype with a modest effect on viability. We demonstrate that *oskar* mRNA interacts with Rab7 suggesting possible role for the Mon1-Rab7 axis in the transport of *oskar*. We show that Mon1 in the PFCs, regulates PIP_2_ levels to influence accumulation of Par-1 in the oocyte. We propose that Mon1 regulates *oskar* localization in two distinct ways: cell autonomously in the germline by regulating Rab7, and non-cell autonomously through the PFCs by regulating accumulation of Par-1.

**Summary statement:** Mon1, an established Rab converter, has roles in embryonic axial patterning, modulating transport, localisation and anchoring of posteriorly localised mRNA during oocyte maturation. Mon1 influence is both cell autonomous, from within the oocyte and non-cell autonomous, through posterior follicle cells.

## Introduction

Asymmetric localisation of mRNA coupled with translational control is one of the key mechanisms used to establish cell asymmetry in eukaryotes. This is seen in budding yeast where asymmetric transport and localisation of ASH1 mRNA to the daughter cell is essential for mating type switch in daughter cell (Long *et al*. 1997; Singer-Kruger and Jansen 2014). In polarised cells like neurons, mRNAs are transported along axons and stored at the synapses as ribonucleoparticles to be translated locally when required (Jung *et al*. 2012).

In *Drosophila*, establishment of the embryonic anterior-posterior (A-P) axis is dependent on asymmetric localization of *bicoid* (*bcd)* and *oskar* (*osk*) mRNAs to the anterior and posterior poles of the oocyte respectively (Driever and Nusslein-Volhard 1988; Kim-Ha *et al*. 1991). Localization of these mRNA requires the RNA binding protein Staufen (*Stau*; (St JOHNSTON *et al*. 1991; Ferrandon *et al*. 1994)). In case of *osk*, Staufen is also required for its translation at the posterior. Staufen has five double-stranded RNA binding domains (dsRBDs) of which domain 2 is required for posterior localization of *osk*; domain 3 binds dsRNA in vitro and is required for localization of both *osk* and *bcd* mRNAs; domain 5 alleviates repression of the transcript to enable translation of *osk* mRNA (St Johnston and Nusslein-Volhard 1992; Micklem *et al*. 2000; Ramos *et al*. 2000).

From several studies, it is well established that the localisation of *osk* mRNA is microtubule-dependent and involves kinesin motors (Brendza *et al*. 2000; Cha *et al*. 2002; Lu *et al*. 2020). The plus-end of the microtubules orient themselves to the posterior in response to an as yet unknown signal emanating from the posterior follicle cells (PFCs) (Gonzalez-Reyes and St Johnston 1994). One of the earliest detectable events downstream of this signal is di-phosphorylation of myosin regulatory light chain (MRLC), followed by accumulation of Par-1 (Doerflinger *et al*. 2010; Doerflinger *et al*. 2022).

Upon reaching the posterior, *osk* mRNA is translated from two alternate start codons leading to synthesis of both long and short isoforms of the protein (Ephrussi and Lehmann 1992; Markussen *et al*. 1995; Vanzo and Ephrussi 2002). These isoforms perform distinct functions: Long Osk (L-Osk) regulates actin and short Osk (S-Osk) mediates assembly of the pole plasm (Markussen *et al*. 1995; Vanzo and Ephrussi 2002).

Endocytic proteins appear to play an important role in localizing *osk* mRNA (DOLLAR *et al*. 2002; Tanaka and Nakamura 2008; Tanaka *et al*. 2011; Tanaka and Nakamura 2011; Tanaka *et al*. 2021). Further, L-Osk itself localises to endosomes and triggers endocytosis at the posterior which is essential for anchoring *osk* mRNA (Vanzo *et al*. 2007). However, despite these studies, a clear understanding of the interplay between the endocytic pathway and the molecular mechanisms leading to localization and anchoring of mRNAs is lacking.

Monensin sensitivity 1 (Mon1) is an evolutionarily conserved protein which complexes with CCZ1 (MC1 complex) and functions as a guanine nucleotide exchange factor (GEF) for Rab7 leading to its activation and facilitating the conversion of an early endosome to a late endosome. In *Drosophila*, loss of *mon1* results in enlarged Rab5-positive early endosomes; the mutants have a short lifespan, motor deficits, and exhibit non-sex specific sterility, all of which can be rescued by driving expression of *mon1* in only in the small subset of octopaminergic-tyraminergic neurons using *tdc2-GAL4* (Yousefian *et al*. 2013; Deivasigamani *et al*. 2015; Dhiman *et al*. 2019).

In this study we identify a role for Mon1 in A-P patterning. Loss of maternal *mon1* leads to mislocalization and loss of anchoring of *osk* and *bcd* mRNAs in the embryo. In mutant oocytes Staufen appears clumped, the level of Osk and Par-1 proteins is low and the ooplasm shows presence of abnormal actin rings. We find that restoring Mon1 function in the germline suppresses the Staufen and Osk phenotypes and rescues lethality. Further, RNA immunoprecipitation experiments show that Rab7 interacts with *osk* mRNA suggesting a role for late endosomes in the transport of *oskar*.

In contrast, expressing Mon1 in the PFCs leads to a significant suppression of the Par-1 phenotype with a modest rescue of lethality indicating that the regulation of Par-1 accumulation is largely through the PFCs. Our results suggest this regulation to be mediated through modulation of PIP_2_. Supporting this, we find that knockdown of PIP4K in the PFCs, affects accumulation of Par-1 in the oocyte.

Collectively, our results identify two distinct roles for *mon1*. In the germline the Mon1-Rab7 axis regulates Staufen and localization of *osk* mRNA. In the PFCs, Mon1 regulates PIP_2_ which regulates accumulation of Par-1 in the oocyte.

## Results

### Loss of maternal *mon1* leads to defects in embryonic patterning

Homozygous *mon1*^Δ^*^181^* mutants (*mon1*^Δ^*^181/^*^Δ^*^181^*) have a short life span, exhibit motor defects and non-sex-specific sterility (Deivasigamani *et al*. 2015; Dhiman *et al*. 2019). The sterility in mutant females is caused by a failure of the egg chambers to undergo vitellogenesis due to downregulated expression of insulin-like peptides, leading to impaired insulin signaling. Interestingly, we found that expressing *mon1* in *tdc2*-positive octopaminergic-tyraminergic neurons (OPNs/OANs) alone, is sufficient to rescue both, lethality and sterility and the ‘rescued’ mutants (*mon1*^Δ^*^181/^*^Δ^*^181^*, *tdc2-GAL4; UAS-mon1:HA*) can lay eggs at rates comparable to wildtype animals (Dhiman *et al*. 2019).

Intriguingly, we found that less than 5 % of these eggs hatch into larvae. Moreover, the lethality was greater than 95% irrespective of the male genotype (Fig. 1B). In contrast, eggs laid by wildtype (*w^1118^*) females crossed with wildtype (*w^1118^*) or ‘rescued’ mutant males (*mon1*^Δ^*^181/^*^Δ^*^181^*, *tdc2-GAL4; UAS-mon1:HA*) were found to hatch normally (Fig. 1B). Together, this suggested a maternal requirement of *mon1* for normal development. In all subsequent sections of the manuscript, we refer to *mon1*^Δ^*^181/^*^Δ^*^181^*, *tdc2-GAL4; UAS-mon1:HA* animals as *mon1^m^* since they are germline ‘mutant’ for *mon1* and refer to eggs laid by *mon1^m^* females as *mon1^m^* embryos.

**Figure 1.**
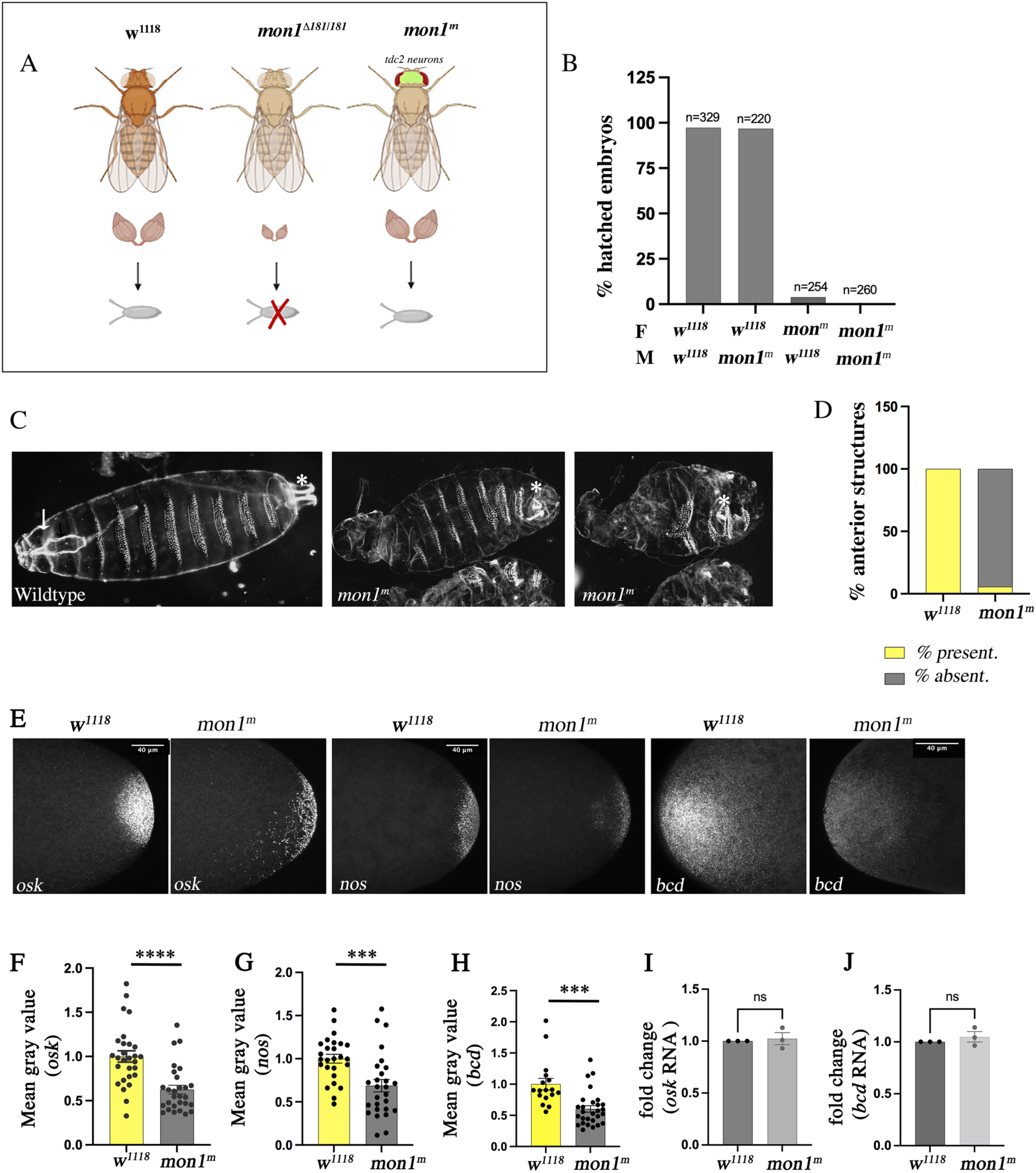
*Mon1^m^* embryos exhibit A-P patterning defects and mislocalization of *osk* and *bcd* mRNAs. **(A)** Schematic describing the genotype and phenotype of *mon1^m^* (*mon1*^Δ^*^181/^*^Δ^*^181^*, *tdc2-GAL4; UAS-mon1:HA*) mutants. *Mon1*^Δ^*^181/^*^Δ^*^181^* do not produce eggs. Expression of *UAS*-*mon1:HA* in *tdc2 postive octopaminergic-tyraminergic* neurons in *mon1^m^* (represented as a green patch in the adult brain) rescues egg production. **(B)** Maternal e>ect of *mon1*. Eggs laid by *mon1^m^* mothers fail to hatch. (*mon1^m^* X *w^1118^*: 3.14% hatching; *mon1^m^* X *mon1^m^*: 0.38% hatching) **(C-D)** Representative images of cuticle preps of *wild-type* and *mon1^m^* embryos. Anterior is to the left and posterior to the right. The anterior and posterior structures are indicated by an arrow and asterisk (*) respectively. More than 90% *mon1^m^* lack anterior structures. (**E**) Fluorescence mRNA *in-situ* against *osk*, *nos* and *bcd* mRNAs. The mRNAs show a more di>use pattern of localisation in the mutants. **(F-H)** *mon1^m^* embryos show reduced fluorescence intensity for *osk*, *nos* and *bcd* mRNAs at the poles. **(I-J)** Level of *osk* and *bcd* mRNA are unaltered in mutant embryos.

We performed a cuticle analysis to determine the cause of lethality in *mon1^m^* embryos. Interestingly we found a complete absence of head structures, with 30% of embryos (n=178) also lacking posterior structures. This was accompanied by mild ventralization and fused denticle belts, suggesting defects in A-P patterning (Fig. 1C, D). These defects were not seen in embryos derived from w*^1118^* females crossed to *mon1^m^* males.

A-P patterning is dictated by localization of *bicoid* (*bcd)* mRNA at the anterior pole, and *oskar* (*osk*) and *nanos* (*nos*) mRNAs at the posterior end. We therefore performed RNA in situs on 0-1hr *w^1118^* and *mon1^m^* embryos to check if localization of these mRNAs was altered. In wildtype embryos, the bright intense staining for *bcd*, *osk* and *nos* mRNAs was seen localized to a small region at their respective terminals. Interestingly, in *mon1^m^* embryos localization of all three transcripts was strongly affected. In most cases even though the transcripts localized to the correct pole, the staining appeared weak and diffuse (Fig.1E).

We quantified this phenotype by measuring the staining intensity within a defined area at the two poles and found a significant decrease in the intensity of all three transcripts in *mon1^m^* embryos (Fig. 1F-H). Interestingly, embryos with mislocalized *osk* and *nos* mRNA showed presence of pole cells at these ectopic locations indicating that the ability to assemble germplasm was not compromised in these mutant embryos.

To determine whether the decrease in staining intensity of the RNA was due to a change in transcript levels, we performed quantitative RT-PCR for *bcd* and *osk* using mRNA isolated from *mon1^m^* ovaries. No significant difference was seen in transcript levels (Fig. 1I-J). This indicated that the decrease in staining intensity was likely to be caused by poor accumulation of mRNA due defects in transport and/or anchoring of these transcripts at their respective poles. In this study we have focused our attention on the role of *mon1* in regulating localization of *osk*.

### Staufen form and transport is perturbed in *mon1^m^* egg chambers

Transport and localization of *osk* mRNA to the posterior takes place during oogenesis. This is mediated by Staufen (Stau), a key RNA binding protein that is also required for translation of *osk* at posterior (St JOHNSTON *et al*. 1991; Micklem *et al*. 2000; Ramos *et al*. 2000). The strong correlation between localization of Staufen and *osk* mRNA has led many studies in the field to use Staufen as a proxy for *osk* localization.

Given the importance of Staufen in localization of *osk*, we examined *mon1^m^* egg chambers for change in expression and localization of the protein, if any. We staged egg chambers based on previous descriptions, in which presence of border cells in the anterior follicular epithelium marks stage 8; delamination of these cells into the nurse cell region marks the beginning of stage 9, which ends when these cells reach the interface between the nurse cells and the anterior end of the egg chamber (Denef and Schupbach 2003). Accordingly, we have categorized egg chambers (Schematic, Fig. 2A) with border cells positioned between the anterior tip and midpoint of the nurse cell region as early stage 9 (E9) while those with cells placed between the midpoint and anterior end of the oocyte as late stage 9 (L9; Fig. 2A).

**Figure 2.**
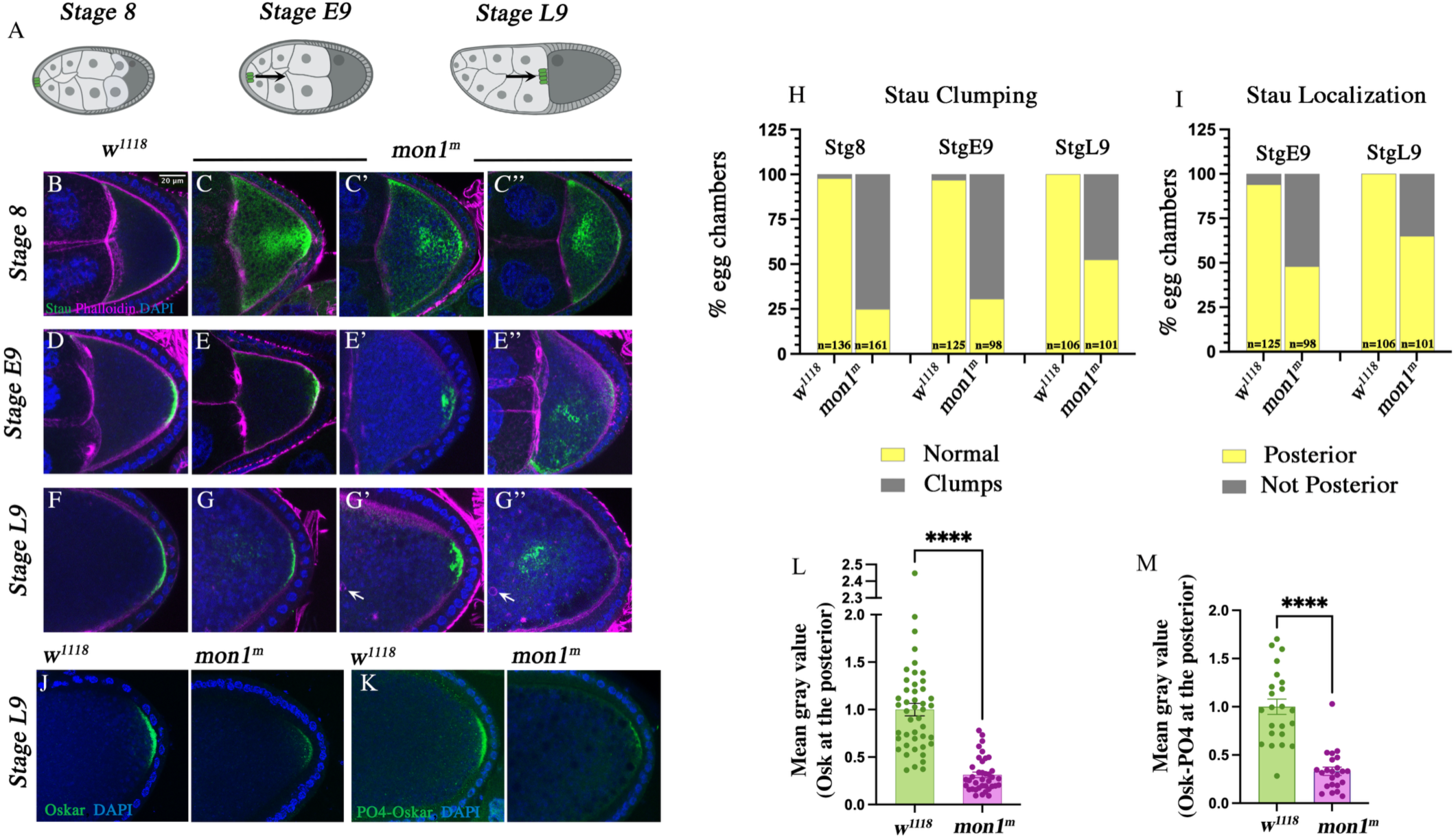
*Mon1^m^* oocytes show clumping and delay in localization of Staufen to the posterior. **(A)** A schematic showing staging of egg chambers for characterization of Stau localization. At stage 8, the anterior polar cells (green) are positioned in the follicle cell layer. Egg chambers with anterior polar cells (green) positioned between the anterior follicle cells and midpoint of nurse cell region (arrow) were scored as stage early 9 (E9); those with polar cells positioned between the midpoint and anterior of the oocyte as late 9 (L9) (**B-G)** Wildtype and *mon1^m^* egg chambers at stages 8, E9 and L9 stained with Anti-Stau (green), phalloidin (magenta) and DAPI. Arrows in G’ and G’’ point to actin rings seen in mutant oocytes. **(H-I)** Quantification of the Stau phenotype in wildtype and *mon1^m^* mutant egg chambers. **(J-K)** Wild type and *mon1^m^* egg chambers stage at L9 stained for Osk (**J**) phosphorylated Oskar (K). **(L-M)** Staining intensity of Oskar (**L**) and phosphorylated Oskar (**M**) is reduced (∼67%) in *mon1^m^* (L: Control (n= 45), *mon1^m^* (n=38); M: Control (n=23), *mon1^m^* (n=23)). **** indicates p<0.0001.

In wildtype stage 7 egg chambers, Staufen staining appears distributed throughout the oocyte. At stage 8, a crescent-like staining is seen at the posterior cortex in most egg chambers (Fig. 2B), although in some, staining is also seen distributed in the oocyte reflecting the on-going process of posterior localization. Post stage E9, all oocytes show a clear crescent-like staining at the posterior cortex (Fig. 2D, F, I).

Interestingly, Staufen staining in *mon1^m^* oocytes appeared as amorphous ‘clumps’ and at times as a ‘diffuse cloud’ with the latter being reminiscent of phenotypes reported in mutants affecting the cytoskeleton (Doerflinger *et al*. 2022). The Staufen phenotype was both striking and highly penetrant with approximately 75% of stage 8 and 69% of E9 oocytes displaying this unusual staining pattern (Fig. 2C, E, H). At stage L9, clumping could be clearly discerned in about half the oocytes (47%); in others the staining appeared faint or undetectable at the posterior. In such cases, Staufen was found frequently mislocalized to middle of the oocyte or close to the posterior cortex (Fig. 2G, H).

In addition to clumping, we also observed a mild delay in Staufen localization. In wildtype ovaries, nearly all stage E9 oocytes show Staufen staining at the posterior cortex. In comparison, only 52% of mutant oocytes showed a distinct crescent (Fig. 2E, I). In the remaining 48%, Staufen localized partially to the posterior or remained dispersed. Staining was detected in the anterolateral regions, indicating defects in the transport process. The clumping and delay phenotypes were also observed in oocytes derived from heteroallelic mutant females (ie. *mon1*^Δ^*^181/^* ^Δ^*^15C,^ elav-GAL4>UAS-mon1HA*), thus ruling out allele specific effects and confirming that the phenotypes are indeed due to loss of maternal *mon1*.

Since Staufen is also required for translation of *osk* mRNA at the posterior (Micklem *et al*. 2000), we checked whether the change in form and localization of Staufen impacts *osk* translation. Indeed, a significant decrease in Osk staining was observed in the mutants (Fig. 2J). Further, oocytes with mislocalized Staufen, also showed ectopic translation of *osk,* indicating loss of repression and translational control. To evaluate the extent of decrease, we measured the intensity of Osk staining in stage 9L oocytes and found a 67% decrease in staining intensity in the mutants (Fig. 2L).

*Osk* mRNA produces two independent isoforms of the protein through selective use of alternate start codons (Markussen *et al*. 1995). Long Oskar (L-Osk) regulates actin while the short isoform (S-Osk) controls germ plasm assembly and therefore the number of future primordial germ cells. The level of S-Osk is tightly controlled through a process involving phosphorylation followed by degradation through the proteasomal system (Morais-De-Sa *et al*. 2013). Sequential phosphorylation by Par-1 followed by GSK3β/Shaggy generates a phosphor-degron that targets Osk for degradation (Morais-De-Sa *et al*. 2013).

We assessed whether Osk in *mon1^m^* oocytes undergo enhanced degradation, by staining the ovaries with an antibody against phosphorylated Osk (S248;(Morais-De-Sa *et al*. 2013)) and found reduced phospho-Osk staining in the mutants (Fig. 2K). The decrease in intensity was comparable to unphosphorylated Osk (Fig. 2M) suggesting that increased degradation is unlikely to be the reason for low Oskar levels in the mutants. These results, together with the fact that transcript levels are unaltered (Fig. 1I), suggests that defects in RNA localization caused by transport and/or anchoring is likely to be the cause for decrease in Osk levels in the mutants although one cannot completely rule out translational defects as well.

### Long Osk and Par-1 levels are reduced in *Mon1^m^* oocytes

A consistent feature observed in *mon1^m^* oocytes was the presence of actin rings in the ooplasm (Fig. 1G’-G’’). These rings are most prominently visible stage L9 onwards and we found that their presence could be used to unequivocally distinguish wildtype and mutant egg chambers. As L-Osk regulates actin, we stained *mon1^m^* egg chambers for L-Osk to determine whether levels of this isoform are altered in mutants. Interestingly, we not only observed a decrease in staining intensity, but also mis-localization of the protein along the posterior and lateral regions of the cortex (Fig. 3B). Staining could also be detected in vesicular structures budding off the cortical membrane. This was unexpected since no such pattern was observed with the Osk antibody (Fig. 2J, K) which detects the C-terminal region common to both isoforms suggesting that the common epitope is either lacking or not recognized by the antibody.

**Figure 3.**
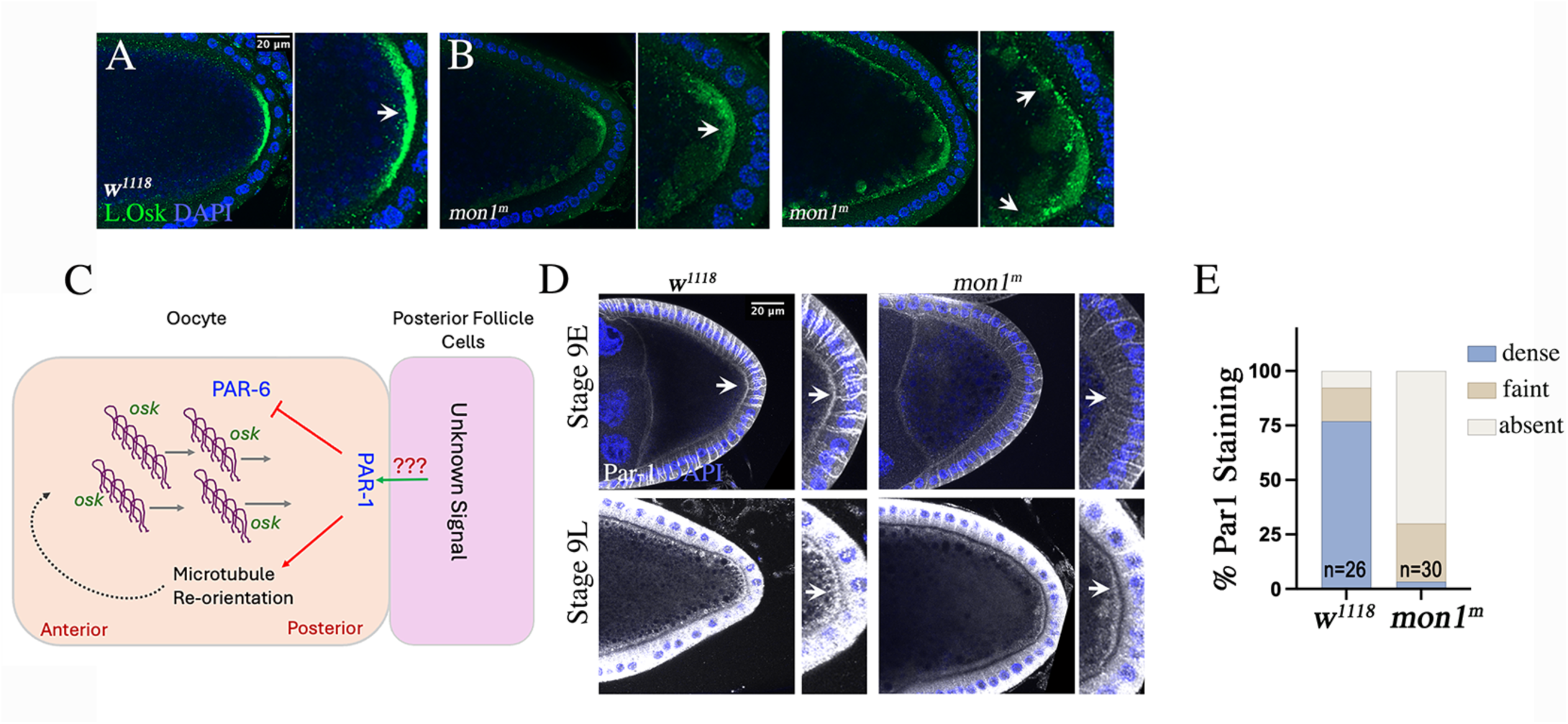
Localization of Long Osk and Par-1 is perturbed in *mon1^m^* mutants. **(A-B)** Stage L9 oocyte stained for Long Oskar (green). Staining intensity is reduced in *mon1^m^* oocytes. Mis-localization is seen along the lateral cortex and in vesicular structures (arrow). **(C)** Schematic describing regulation of *osk* localization by Par1. **(D)** Par1 staining in stage E9 and stage L9 egg chambers. A crescent-like staining (arrows) is seen at the posterior. *Mon1^m^* oocytes show reduced staining (arrow). **(E)** Quantitation of the Par-1 phenotype in stage E9 oocytes: 70 % of mutant oocytes lack Par-1 staining compared with 8% in controls.

Transport of *osk* to the posterior is dependent on microtubules. Par-1 is a serine-threonine kinase that functions downstream of an as yet unknown signal coming from the posterior follicle cells to regulate re-orientation of microtubules at stage 7 (Fig. 3C; (Shulman *et al*. 2000; Tomancak *et al*. 2000; Doerflinger *et al*. 2006; Doerflinger *et al*. 2010)). We therefore examined egg chambers at stages E9 and L9 to determine if Par-1 levels are altered in *mon1^m^* mutants and found reduced Par-1 staining at both E9 and L9 stages (Fig. 3D). We scored the phenotype in stage E9 oocytes by categorizing the staining into dense, faint or absent. Interestingly, 70% of mutant oocytes showed absence of Par-1 staining (Fig. 3E). Consistent with this, we also observed defects in the microtubule network (data not shown). Based on the hierarchy of events involved, this result suggests an upstream role for Mon1 in localization of *osk*.

### Mon1 is expressed in the oocyte and enriched in anterior and posterior polar cells

Next, we sought to determine the expression pattern of Mon1 in the ovaries. For this we tagged endogenous Mon1 by inserting superfolder (sf) GFP at its genomic locus using CRISPR (*mon1:sfGFP*) and examined expression of GFP using immunostaining. GFP staining was detected in both, the germline and follicle cells. In the oocyte, expression was more clearly visible at early stages and observed along the cortex; in the follicle cell layer, expression was enriched in the posterior polar cells (PPCs; Fig. 4A, A’). Expression in these cells was spatio-temporally regulated with strong expression at stage 8 that decreased with increasing oocyte maturity (Fig. 4A-B) The expression pattern also correlated with the morphogenesis of these two cells: at stage 8 the PPCs appeared spindle like with the proximal tip extending towards the posterior cortex of the oocyte (Fig. 4A, A’); at stage L9 and 10, these cells appeared retracted and more spherical in shape (Fig. 4B, B’). At stage 8 we also detected a ‘platform-like’ GFP staining at the posterior cortex (Fig.4B, yellow arrow) that persisted through till stage L9 (Fig.4B). Interestingly, Staufen localization is sandwiched between this ‘platform’ and posterior cortex (Supplementary Figure. 1).

**Figure 4.**
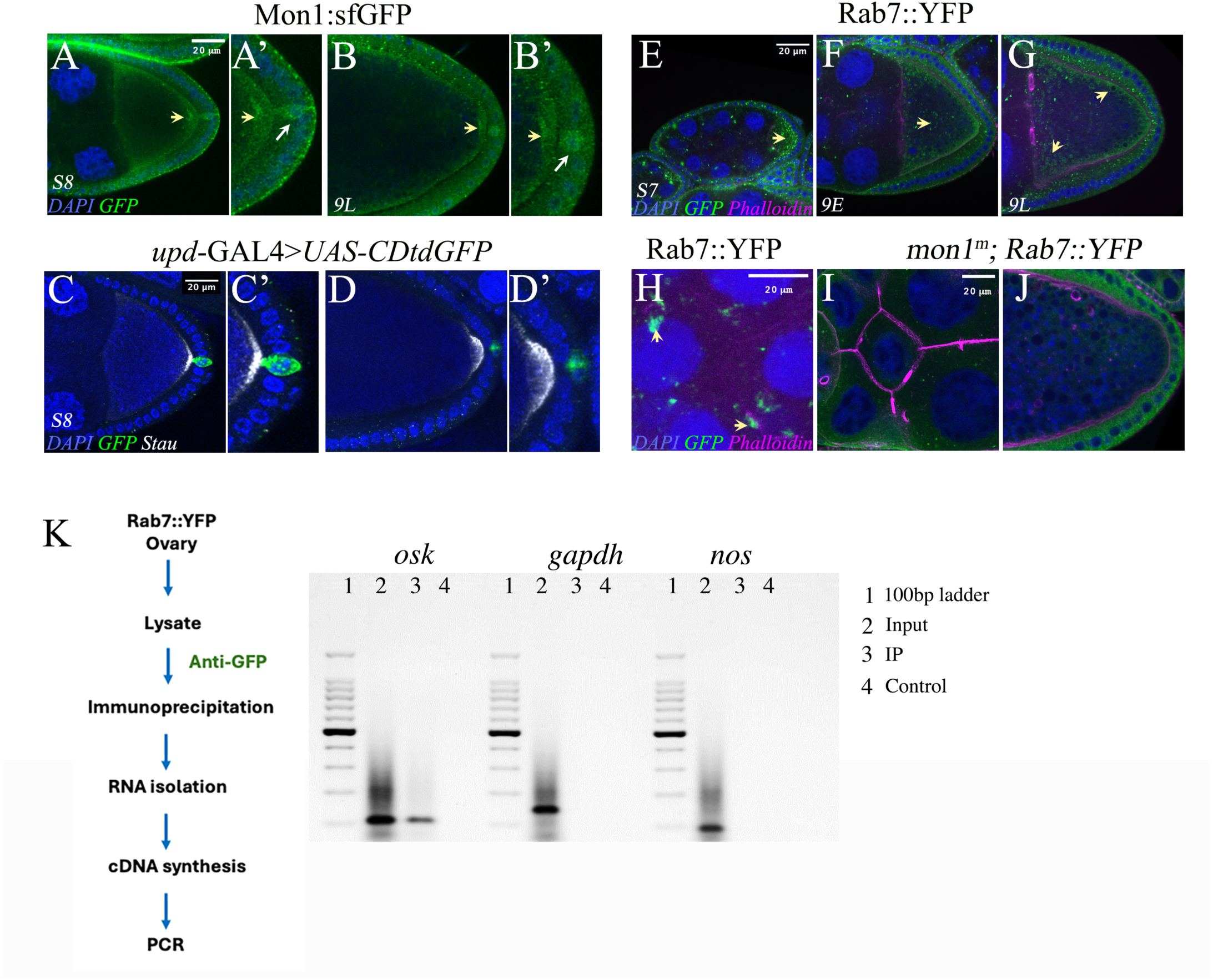
Expression pattern of Mon1 and Rab7 in developing egg chambers. **((A-C)** Mon1::sfGFP expression in egg chambers at stage 8 (A) to stage L9 (B). In addition to the oocyte, enriched temporal expression is seen in PPCs. **(C-D)** *upd*-GAL4>*UAS-CD4tdGFP* egg chambers stained for GFP (green) and Stau (white). At stage 8 (C-C’) PPCs (green) are spindle like with an extension towards the cortex. At stage L9, the cells retract and become spherical (D-D’). **(E-G)** Expression of Rab7::YFP in developing oocytes. Density of late endosomes reduces with increasing maturity (F, G., arrows). Staining around vesicles is seen in L9 egg chambers (G, arrows). **(H-J)** Rab7::YFP expression in nurse cells at stage L9. Endosomes concentrate around ring canals (H); they are absent in *mon1^m^* (I-J). **(K)** RNA immunoprecipitation (IP) of Rab7::YFP shows specific association of *osk* mRNA with Rab7. A detailed IP protocol is described in Materials & Methods, and lane/loading definitions are on the right-hand side. For lane 4 (mock control), the antibody was not added to the lysate.

We confirmed that PPCs undergo morphogenesis, by driving expression of *UAS-mCD8GFP* in these cells using *upd-*GAL4. At stage 8, the PPCs are indeed spindle shaped and retract to assume a more spherical morphology after stage 9 (Fig. 4C-D). We also observed that like Mon1::sfGFP, expression of the GAL4 is strong in early egg chambers and decreases with increasing maturity of the oocyte (Fig. 4C-D).

Next, given the role of Mon1 as an activator of Rab7, we examined the expression pattern of Rab7 in egg chambers using a Rab7::YFP line in which Rab7 is tagged at its endogenous locus with myc and YFP at the N and C-terminal respectively. Interestingly we observed numerous Rab7::YFP positive punctae in early previtellogenic oocytes which reduced dramatically at later stages suggesting an important role for Rab7 in early stage egg chambers (Fig. 4E-G). Further, as reported previously (Yu *et al*. 2024), large Rab7::YFP positive vesicles were seen at cortex of more mature oocytes (Fig. 4G). In the nurse cells the staining showed large Rab7 punctae to be frequently present around ring canals (Fig. 4H). As expected, we did not detect these puncta in *mon1^m^* egg chambers (Fig. 4I-J).

Studies in fungi have linked endosomes to RNA transport (Muntjes *et al*. 2021; Schuhmacher *et al*. 2023). In addition, late endosomes and lysosomes have also been identified as platforms for transport and translation of RNA in axons (Cioni *et al*. 2019; Liao *et al*. 2019). Since the abundance of Rab7 staining in stage 7 oocytes coincides with the initiation of events leading to posterior localization of *osk* mRNA, we sought to explore whether Rab7 might be involved in the transport and localization of *osk* mRNA. As a first step, we checked whether Rab7 interacts with *osk* mRNA and to this end, carried out RNA immunoprecipitation (RIP) experiments using ovary lysates from Rab7::YFP animals. Significantly, PCR performed on cDNA generated from immunoprecipitated mRNA showed amplification for *osk* but not for *nos* or *gapdh* indicating specificity of the interaction (Fig.4 K) demonstrating for the first time an interaction between Rab7 and *osk* mRNA. Even though the interaction is likely to be an indirect one, the results nonetheless suggest a role for late endosomes in transporting *osk* mRNA.

### Germline expression of *mon1*::HA rescues the *staufen* and *osk* phenotype

Next, to assess the individual contributions of the germline and somatic follicle cells to the *mon1^m^* phenotype, we performed tissue specific knockdown of *mon1* using *RNAi*. As in the mutants, expression of *UAS-mon1RNAi* (referred to as *mon1i*) using the germline specific *nos-*GAL4 led to clumping of Stau which was most evident at stage 8 where nearly 50% of oocytes showed this phenotype (Fig. 5A, B). The effect was less pronounced at stage E9. Knockdown of *rab7* had stronger effect with nearly 67% and 33% of stage 8 and E9 oocytes showing clumpy Stau (Fig. 5A B). In both cases, we failed to detect the ‘delay’ phenotype observed in the mutants. We also examined Osk levels in these animals and found the staining intensity to be lower in *rab7i*, albeit statistically insignificant.

**Figure 5.**
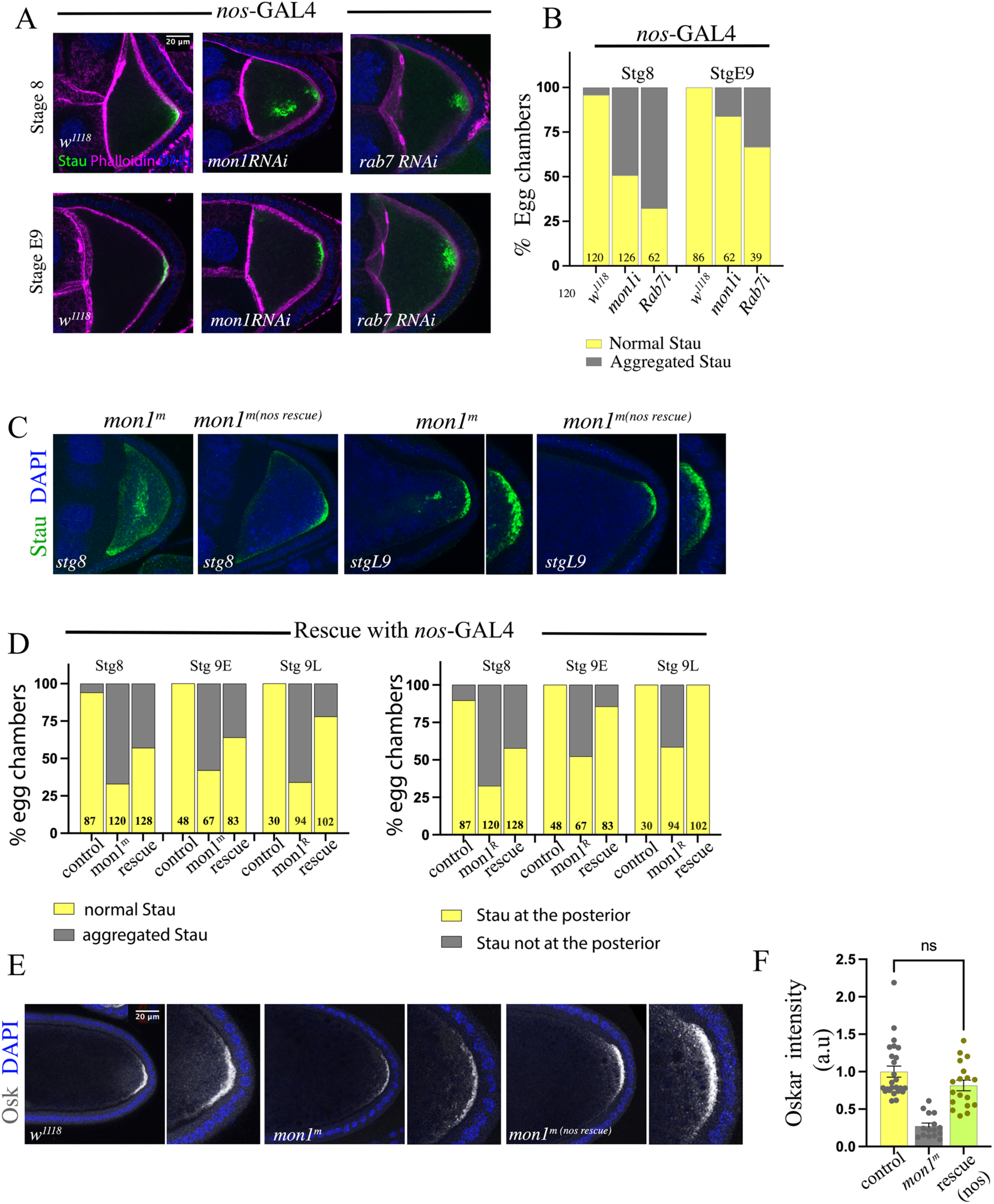
Germline expression of *mon1* rescues Staufen and Osk phenotypes. **(A-B)** Knockdown of *mon1* and *rab7* in the germline, using *nos-Gal4*, leads to clumping of Staufen. **(C-D)** Germline expression of *UAS-mon1HA* in *mon1^m^* suppresses the clumping phenotype by ∼25-30% across all stages. The delay in Staufen localization is also rescued significantly at stage E9 (85%) and L9 (100%) **(E-F)** Germline rescue of *mon1^m^ oocyte* restores Osk levels to near wild type.

We next checked if restoring Mon1 function in the germline rescues the *mon1^m^* phenotype. To do this, we placed *nos*-GAL4 in the background of *mon1^m^*. These animals, hereafter referred to as *mon1^m (nos rescue^*^)^, showed punctate Rab7 staining in the oocyte and nurse cells indicating functional restoration of Mon1. Remarkably, expression of Mon1 in the germline alone was sufficient to rescue many of the *mon1^m^* phenotypes. The Stau phenotype was suppressed by approximately 25-30% across all stages (Fig. 5C, D) and in the others, the phenotype appeared less severe than the mutant. The delay in localization was also rescued, with 85% of stage E9 oocytes showing posterior localization of Stau and 100% at stage L9 (Fig. 5C, D). Finally, Osk expression in these oocytes was comparable to wildtype (Fig. 5 E-F). Notably, the abnormal actin rings in the mutant were completely absent (Supplementary Figure. 2). Taken together, these results, indicate a crucial germline role for Mon1 in regulating *osk* localisation.

### *Mon1* in posterior follicle cells regulates the localization of Par-1 at the posterior

The PFCs play a crucial role in regulating polarity in the oocyte by releasing an as yet unknown signal at stage 7 leading to accumulation of Par-1 and reorientation of microtubules (Gonzalez-Reyes and St Johnston 1994). The differentiation of PFCs is regulated by JAK-STAT signaling and contact between these cells and the oocyte is essential for localization and accumulation of Par-1 and *osk* mRNA (Assa-Kunik *et al*. 2007; Mallart *et al*. 2024).

To explore the role of *mon1* in the PFCs, we used *c306-GAL4* to knockdown expression of the gene using RNAi. Interestingly, Stau in these oocytes appeared clumpy similar to *mon1^m^*. The phenotype was observed in approximately 60% and 66% of stage 8 and E9 oocytes respectively (Fig. 6A, B). Further, Osk staining was also reduced by 55% (Fig. 6C, D) and intensity of Par-1 staining was much reduced (Fig. 6E, F), with 37% of stage E9 oocytes lacking staining.

**Figure 6.**
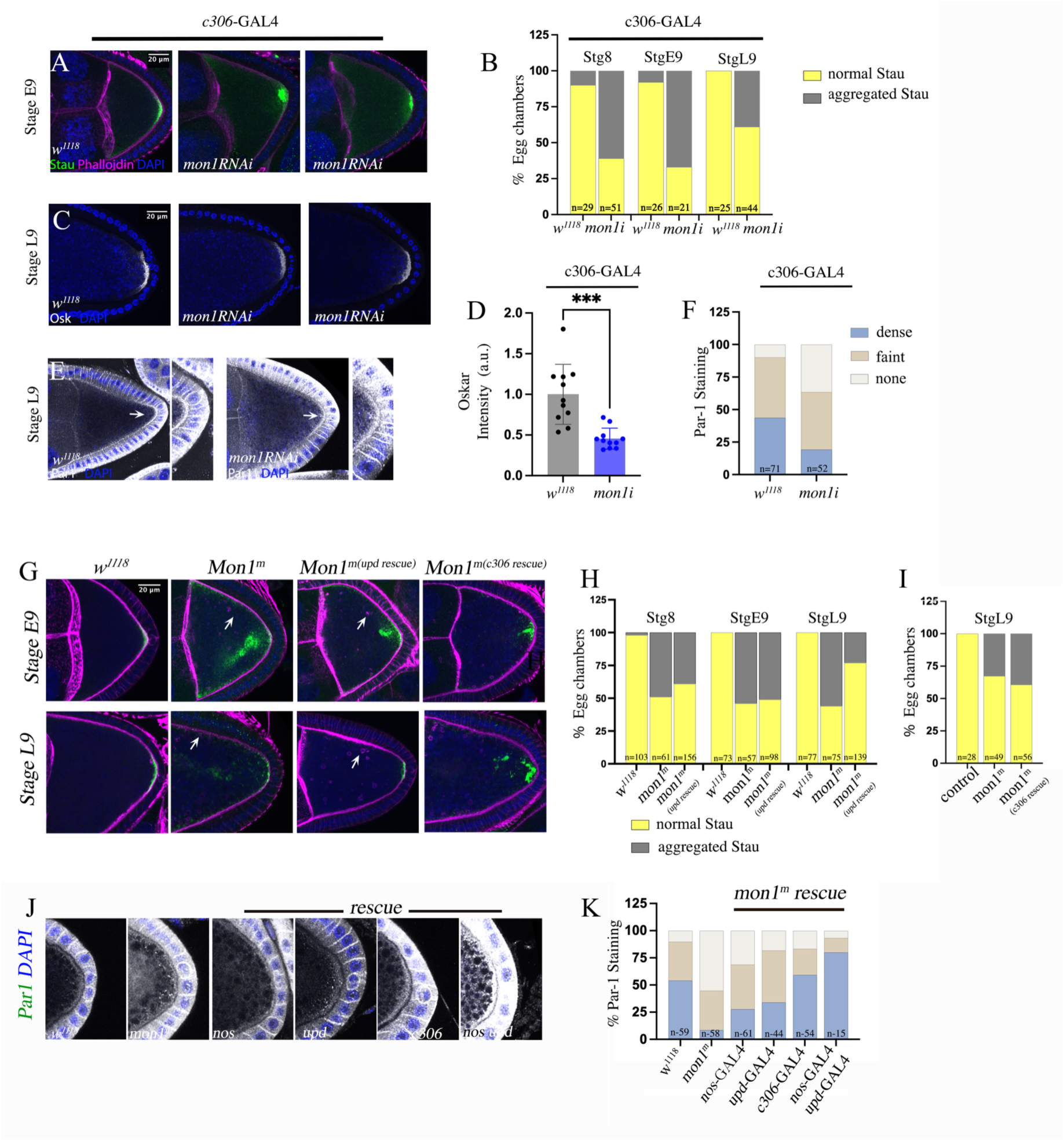
Mon1 in posterior follicle cells regulates Par1 accumulation at the posterior cortex. **(A-B)** Knock-down of *mon1* in posterior follicle cells using *c306-Gal4* leads to aggregation of Staufen (B). **(C-D)** Stage L9 egg chamber. Knockdown of *mon1* in posterior follicle cells leads to decrease in Osk staining at the posterior. (**E-F**) Par1 accumulation at the posterior cortex is reduced in *c306>mon1RNAi oocytes.* 36% of egg chambes lack Par-1 staining. **(G-I)** Expression of *UAS*-*mon1:HA* in PFCs using *upd*-Gal4 and *c306*-Gal4 in *mon1^m^* fails to rescue the Staufen phenotype. **(J-K)** Accumulation of Par-1 is significantly rescued in *mon1^m(c306rescue)^* compared with *mon1^m(nos rescue)^* and *mon1^m(upd rescue)^* animals. In ‘Double-rescue’ (*mon1^m(nos, upd rescue^)* animals, accumulation of Par-1 is significantly higher compared to controls (I). *** indicates p< 0.001.

Interestingly, driving expression of *UAS-mon1::HA* in the PFCs of *mon1^m^* egg chambers using *c306* and *upd GAL4*s (hereafter referred to as *mon1^m(c30C^* ^rescue)^ and *mon1^m(upd^* ^rescue)^ respectively) failed to rescue the Stau phenotype, although some suppression was seen at stage L9 with *upd*-Gal4, (Fig. 6G-I). The characteristic actin rings seen in *mon1^m^* oocytes, continued to persist in these oocytes (Supplementary Figure. 2). Surprisingly, we observed a substantial rescue of the Par-1 phenotype. Compared with *mon1^m(nos rescue)^*, more than 50% of oocytes from *mon1^m(c30C rescue)^* had Par-1 staining comparable to control oocytes. A double rescue using *nos* and *upd* drivers led to a 75% increase with staining in some egg chambers being even more intense than controls (Fig. 6J, K). Collectively, results from the germline and the PFC rescue experiments indicate a bifurcation of Mon1 function: in the germline, Mon1 plays a major role in regulating the form and localization of Staufen which in turn affects localization of *osk* and its translation. Here, Mon1 also affects actin, important for anchoring the mRNA. In the PFCs, Mon1 functions non-cell autonomously to regulate posterior accumulation of Par-1 in the oocyte.

### PIP_2_ in the PFCs regulates accumulation of Par-1 in the oocyte

Studies from multiple labs indicate that the initial recruitment of Par-1 to the posterior cortex, is dependent on actin filaments (Doerflinger *et al*. 2006) where it promotes polarization of microtubules for posterior localization of *osk* mRNA. The subsequent translation of *osk* triggers a feedback loop in which Par-1 stabilizes Osk protein through phosphorylation, thus enabling it to recruit more Par-1 to the posterior, which in turn, leads to further polarization of microtubules, promoting biased posterior transport of *osk* mRNA (Riechmann *et al*. 2002; Riechmann and Ephrussi 2004). Studies have also shown that contact between posterior follicle cells and the oocyte cortex is required for accumulation of Par-1. It is therefore possible that the contact between PFCs and the oocyte, in addition to mediating signal transduction, also alters local membrane composition that impacts accumulation of Par-1. PIP_2_ is a well-known regulator of actin dynamics. Local increase in PIP_2_ levels positively regulates actin polymerization (Logan and Mandato 2006; Senju and Lappalainen 2019). Based on this we decided to test whether Mon1 might affect accumulation of Par-1 by modulating levels of PIP_2_. Using PH-PLCδ-mcherry as a reporter, we explored whether knockdown of *mon1* leads to changes in PIP_2_ levels. We used *upd*-GAL4 in this experiment because of its restricted expression pattern and ease of analysis. Consistent with our hypothesis, knockdown of *mon1* led to a significant increase in PIP_2_ reporter intensity in the PPCs. We quantified the increase by staining the ovaries with an anti-RFP antibody. Compared with control, we observed many more intensely stained intracellular RFP-positive puncta in PPCs at stage L9 (Fig. 7A). We interpret this result to indicate that downregulating Mon1 leads to a retention of PIP_2_ enriched vesicles in the PPCs. To confirm that the change in intensity reflects lipid levels, we measured intensity of the reporter upon knock-down of PIP4K, a PIP_2_ generating enzyme. As expected, a decrease in reporter intensity was seen in these cells indicating reduced synthesis of PIP_2_ (Fig. 7B).

**Figure 7.**
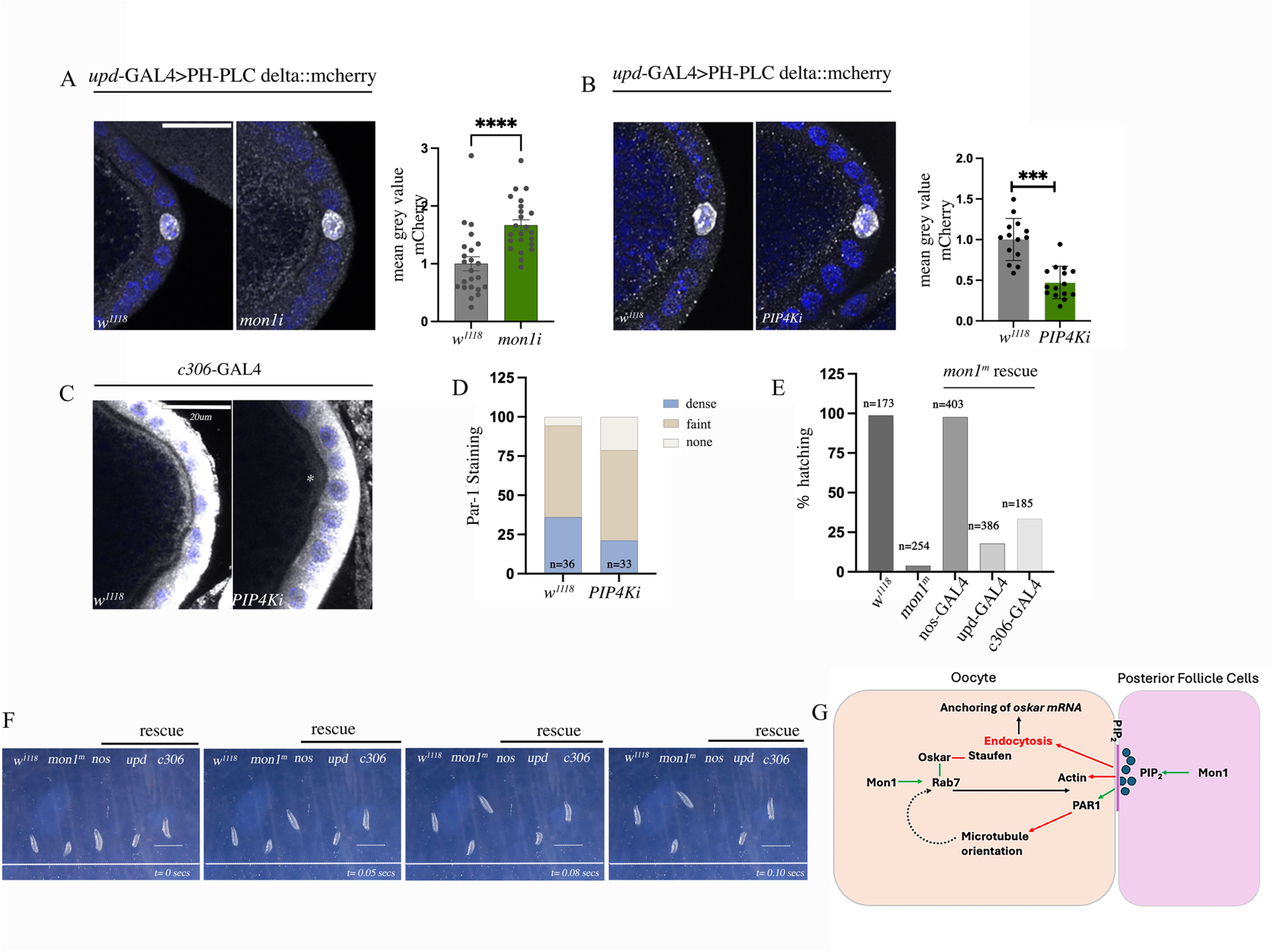
Mon1 regulates Par-1 accumulation through modulation of PIP2. **(A)** Stage L9 oocyte expressing the PIP2 reporter PH-PLCδ-mcherry and stained for mcherry using anti-RFP. Knockdown of *mon1* leads to increase in staining intensity of the reporter. **(B)** Knockdown of PIP4K leads to a decrease in the staining intensity of the PIP2 reporter. **(C-D)** Knockdown of PIP4K in posterior follicle cells a>ects accumulation of Par-1 at the posterior cortex. 21% of *c306-Gal4>UAS-PIP4kRNAi* show absence of Par-1 staining compared to 5% in control animals. **(E)** Viability of eggs laid by *mon1^m(rescue)^* animals: 98% of eggs laid by *mon1^m(nos rescue)^* hatch into first instar larvae compared to 4% in *mon1^m^. mon1^m(upd rescue)^* and *mon1^m(c30C rescue)^* show hatching rates of 18 and 33 percent respectively. **(F)** Sequential frames from a movie depicting the rescued *mon1^m^* larvae. The larval mobility of *mon1^m(nos rescue)^* is comparable to *w^1118^* larvae. The movie is included in supplementary information as Movie S1. **(G)** A model for Mon1 function in patterning the posterior cortex. In the oocyte, Mon1 regulates Rab7 to contribute to transport of *osk* mRNA; in the PFCs, Mon1 regulates Par-1 accumulation through modulation of PIP2.

We next asked if reducing PIP_2_ levels affects accumulation of Par-1 in the oocyte. Interestingly, knockdown of PIP4K in the PFCs led to reduced Par-1 accumulation (Fig. 7D) in the oocyte with 21% oocytes lacking staining compared to 5% in controls (Fig. 7D). Finally, we checked whether rescue via the germline or PFCs restores viability in *mon1^m^* embryos. Remarkably, embryos from *mon1^m(nos rescue)^* mothers showed a near 100% hatching rate, similar to wildtype; a more modest but nonetheless a substantial increase in hatching (∼25%) was seen with *upd* and *c306* driver lines (Fig. 7D). The robustness of the rescue was reflected in the crawling ability of the larvae. *Mon1^m^* larvae are typically sluggish and exhibit poor mobility. In contrast, *mon1^m(nos rescue)^* and *mon1^m(c30C rescue)^* larvae were active with the mobility of the former being comparable to wildtype. In contrast, *mon1^m(upd rescue)^* larvae were less mobile (Fig. 7E; see Movie S1). Together these results indicate that even though *mon1* in the germline is necessary and sufficient to rescue patterning and lethality, it has a redundant but nonetheless significant regulatory role in the PFCs that can rescue patterning and lethality albeit at a lower frequency.

Together, these findings reveal that Mon1 coordinates A-P patterning through both, the germline and PFCs. In the germline the Mon1-Rab7 pathway regulates Osk transport, translation and anchoring, through regulation of Staufen and actin; In the posterior follicle cells, Mon1, through regulation of PIP_2_, functions to prefigure the posterior cortex to facilitate accumulation of Par-1 and promote polarized transport.

## Discussion

Previous studies from the Ephrussi and Nakamura labs strongly suggest a role for the endocytic pathway in regulating localization of *osk* mRNA. L-Osk localizes to endosomes in the oocyte and is thought to recruit endocytic proteins to trigger endocytosis essential for actin remodelling and anchoring of *osk* mRNA and consequently the germplasm (Vanzo *et al*. 2007; Tanaka and Nakamura 2008). Loss of endocytic proteins Rab11 and Mon2, which regulate vesicle recycling and Golgi-ER trafficking respectively, leads to mislocalization and loss of anchorage of Osk (Dollar *et al*. 2002; Tanaka and Nakamura 2011). Furthermore, loss of Rabenosyn-5 (rbsn-5), an effector of Rab5, influences localisation and anchoring of *oskar* mRNA by regulating orientation of microtubules (Tanaka and Nakamura 2008).

In this study we provide the systematic evidence that Mon1-a key regulator of the endo-lysosomal pathway regulates *osk* mRNA localization in the oocyte and demonstrate for the first time, an interaction between Rab7 and *osk* mRNA. We also uncover a role for Mon1 in regulating a PIP_2_ dependent signal from the PFCs that regulates Par-1 localization.

### How does Mon1-PIP_2_ regulate Par-1 accumulation?

Accumulation of Par-1 at the posterior cortex of the oocyte is one of the first steps in patterning of the posterior. We find that *mon1^m^* oocytes fail to accumulate Par-1. This change can be detected at stage E9 indicating that the process is perturbed right from the early stages of oogenesis (Fig. 4). Since the phenotype can be strongly suppressed by expression of the gene in PFCs this indicates that the regulation is largely non-autonomous. We also find that the regulation by Mon1 is mediated by changes in the level of PIP_2_ suggesting a role for this lipid in regulating Par-1 accumulation (Fig. 6, 7).

That PIP_2_ affects Par-1 accumulation is known from studies done on *PIP5K/skittles* mutant oocytes which show a global decrease in PIP_2_ levels in the cortical membrane. Par-1 levels at the posterior is found to be significantly reduced in these mutants (Compagnon *et al*. 2009). Interestingly, localization and accumulation of Par-1 is also dependent on mechanical contact between the PFCs and the oocyte. Disrupting this connection is found to lead to removal of Par-1 and premature accumulation of the Par6/Bazooka complex (Milas *et al*. 2023; Mallart *et al*. 2024).

Intriguingly, the initial accumulation of Par-1 which begins at stage 8, coincides with elevated expression of Mon1 in the PPCs (Fig. 4). At this stage the PPCs together with the surrounding PFCs are in close contact with the posterior cortex of the oocyte (Mallart *et al*. 2024). Based on this one could speculate that *mon1* in the PFCs (including the PPCs) promotes release of PIP_2_ positive vesicles, which, because of the contact between PFCs and the oocyte at stage 8, leads to a ‘local’ increase in the concentration of the lipid at the posterior cortex. This, in turn, could trigger signaling leading to reorganization of actin essential for capture and accumulation of Par-1. Alternately, local increase in PIP_2_ could also trigger endocytosis causing remodeling of actin essential for Par-1 accumulation. Each of these possibilities would need to be tested.

It is also remarkable that driving expression of *mon1* in the PPCs or PFCs of *mon1^m^* (*mon1^m(upd rescue^* and *mon1^m(c30C rescue^)* alone, is sufficient to suppress embryonic lethality of *mon1^m^* albeit only to a modest level. This contrasts with the near complete rescue seen in eggs laid by *mon1^m(nos rescue)^* despite the absence of a strong rescue of Par-1 (Fig. 6, 7). This indicates not only a crucial role for *mon1* in the germline but also suggests that a minimal threshold of Par-1 is sufficient to ensure efficient execution of downstream events leading to *osk* localisation. Furthermore, this points to the cross-talk between the PFCs and oocyte as constituting a redundant mechanism, sufficient to restore patterning and germplasm formation in response to perturbations in the germline.

### Mon1-Rab7 axis and regulation of *osk* mRNA

Another novel and interesting finding to emerge from our study is the association between *osk* mRNA and Rab7 (Fig. 4). The current understanding of *osk* mRNA transport shows that the transcript organizes into membraneless ribonucleoprotein (RNP) complex, presumably with Staufen, and is carried to the posterior by kinesin motors (Brendza *et al*. 2000; Bose *et al*. 2022; Bose *et al*. 2024). However, live imaging studies of *osk* mRNA and Staufen suggest that both molecules are initially trafficked independently and, at a later point, move together towards the posterior (Mhlanga *et al*. 2009). Our result showing an association of *osk* mRNA with Rab7 does not in any way contradict these mechanisms since it is possible that Staufen hitchhikes on late endosomes as is the case with HIV-1 RNA (Muntjes *et al*. 2021). Further, the observation that *osk* mRNA may initially be transported independently of Staufen opens up the possibility that a part of the transport process is on late endosomes.

As late endosomes can also function as platforms for translation, one could speculate that localization of L-Osk to endosomes may begin at the transport step, with mRNA associated with late endosomes being translated primarily into the long isoform. Interestingly, in *mon1^m^* oocytes, the L-Osk antibody stains along the lateral membrane and the same is not detected by anti-Osk that detects the C-terminal common to both L and S-Osk. One explanation for this mismatch is incomplete translation of L-Osk. This would be not only be consistent with the idea of late endosomes being involved in translation, but would also imply that the production of Long and Short Osk is in fact regulated and not stochastic. Indeed, unpublished observations indicate that a 1:4 ratio of L-Osk: S-Osk is required in the oocyte (Vanzo *et al*. 2007). In this scenario, one could also predict that localization of *osk* mRNA from endosomes to the posterior cortex is likely to be Rab11 dependent since Rab11 is associated with trafficking to the plasma membrane. A more detailed study would be needed to address some of these questions. In summary, our study identifies the endo-lysosomal pathway as a crucial regulator of mRNA localization during A-P patterning. Although this study focuses on *osk*, it is likely that similar mechanisms exist to regulate localization of *bicoid* mRNA. This will need to be explored. On a broader note, the study could also have important functional implications for specialized cells like neurons where RNA localization and local translation at the synapse is crucial for functional homeostasis.

## Materials and Methods

### Fly husbandry

All stocks were maintained on regular cornmeal agar medium. *Dmon1*^Δ^*^181^* (*mon1* ^Δ^*^181^*, (Deivasigamani *et al*. 2015)), *UAS-mon1HA* (Thomas Klein lab), *mon1^15C^*(Basargekar et al. 2020); *mon1^m^* refers to *mon1* ^Δ^*^181/^* ^Δ^*^181^; tdc2-GAL4>UAS-mon1HA*, *UAS-Rab7RNAi* (#27051); *UAS-mon1RNAi* (#38600, VDRC), *UAS-PIP4K RNAi* (#65891, BDSC); *nos-Gal4 (#32180,* BDSC), *C306-Gal4* (#3743, BDSC); *upd*-Gal4 (#26796, BDSC); UAS-4c-PLCδ1−PH-mcherry (Basu *et al*. 2020); Rab7::YFP (#62545, BDSC). Standard techniques were used to combine mutants, GAL4 and reporter lines. All RNAi based experiments were conducted at 27°C; rescue experiments were conducted at 25 °C. For all stainings carried out with *mon1^m^* or *mon1* ^Δ^*^181/^* ^Δ^*^15C^,elav-GAL4>UASmon1HA*, mutant females were crossed with *w^1118^* males and kept on yeasted medium for 2-3 days before dissecting the ovaries.

### Generation of *mon1::sf GFP*

Generation of this line was carried out at the flyfacility, cCAMP, BLiSc Campus, Bengaluru. Briefly, CRISPR based homologous recombination strategy was used to generate fly lines expressing Mon1 protein fused at its C-terminus to sfGFP. A suitable gRNA target (CTAAAGCTTAGAATGTGGCATGG) was selected at the stop site of gene mon1. gRNA was cloned in pBFVU6.2 vector (Kondo and Ueda, 2013) using standard PCR based strategy. Homologous recombination (HR) construct was generated by Gibson assembly using pHD-sfGFP-ScarlessDsRed vector (DGRC Stock 1365; https://dgrc.bio.indiana.edu//stock/1365; RRID:DGRC_1365) such that the C-terminus of mon1 protein is in frame with sfGFP. To generate knock-in flies, ∼450 embryos from flies expressing vasa-Cas9 on 3rd chromosome (BDSC#51324) were injected with gRNA (250ng/ul) and HR construct (750ng/ul) using standard microinjection protocol using Eppendorf Femtojet. Surviving adults (400) were self-crossed and F1 progeny were screened for the presence of 3XP3-dsRed marker in the eyes. 33 positive lines were recovered. These were then made into homozygous stocks.

### Immunostaining and Confocal Imaging

Ovaries from 2-3 days old yeast fed females were dissected in 1X PBST(0.2%Tween) at room temperature (RT), fixed with 4%PFA for 20min, followed by 2 PBST washes each for 15 minutes at RT and blocked with PBTX (0.2%Triton-X, 2%BSA) for 40 minutes at 4°C. Staining with primary antibody was carried out overnight at 4°C followed by 4-5 washes at RT with PBTX. Secondary antibody staining was performed overnight at 4°C.

For characterizing expression pattern of Mon1::sfGFP and Rab7::YFP, ovaries are dissected on the ice in 0.2% PBST to preserve endosomal structures and fixed at RT with 4% methanol-free paraformaldehyde for 20 minutes. This was followed by two washes with PBST for 15 minutes each at RT. Blocking was carried out with PBTX (0.2% Triton-X, 2%BSA) for 40 minutes at 4°C and incubated overnight with primary antibody at 4°C.

The following antibodies were used rabbit-anti-Staufen (1:5000, St. Johnston lab), anti-phosphorylated Osk (1:200, St. Johnston lab), guinea pig anti-Osk (1:1500, Tanaka Nakamura lab), rabbit anti-Long Oskar (1:2000, Tanaka Nakamura lab), anti-chicken-anti-GFP (1:200 to 1:500, Thermo Fisher, Cat. no. A10262), DAPI (1:1000, Thermo Fisher, Cat.No. D1306), Alexa Fluor 568 Phalloidin (1:400, Thermo Fisher, Cat no. A12380). The following secondary antibodies were used from Thermo Fisher: anti-rabbit Alexa Fluor 488 (Cat. no. A11034), anti-rabbit Alexa Fluor 568 (Cat no. A110036), anti-chicken Alexa Fluor 488 (Cat no. A11039). All secondary antibodies were used at 1:1000 dilution. Images for qualitative analysis of Staufen and fluorescent in situs were captured using the Leica SP8 confocal system using a 40X oil immersion lens (N.A. =0.95). Images for quantitative analysis of Oskar, Par-1 and PIP_2_ were captured on Zeiss LSM 900 using a 40X oil immersion lens (N.A.=0.95). Vectashield (Vector Labs) was used as the mounting medium.

### RNA in-situ

RNA *in situ* hybridization was carried out on 1-2 hr embryos derived from wildtype (*w^1118^*) and *mon1^m^* females as described in (Ratnaparkhi and Zinn 2007). Digoxigenin labelled sense and antisense RNA probes were generated using full length cDNAs for *oskar* and *nanos*; Probes for *bicoid* were generated using 1.3 kb coding region PCR amplified and cloned into pGEMT-eazy using the following primers: *bcd F* : 5’ TTCAGTTGCCGCCACAATTC 3’ and *bcd R* 5’ CATCGTCGCAGCTCTTGTCC 3’

### Intensity Analysis

Intensity measurement for immunostaining was carried out on sum slice projections by outlining the stained region and measuring the mean fluorescence intensity; intensity of region outside the tissue was subtracted as background noise to obtain the signal value.

For the RNA in situs, a region of 3500-4000μm^2^ outlined at the anterior (*bcd*) and posterior tip (*osk* and *nos*) of the embryo was used measure fluorescence intensity.

### Quantitative RT-PCR

Estimation of *osk* mRNA was carried out using quantitative real-time PCR. Ovaries from 2-3 days old females fed on yeasted medium were dissected and stored in Trizol reagent. RNA isolation followed by cDNA synthesis was carried out according to the manufacturer’s instructions. Approximately 1 µg of total RNA was used for the reverse transcription reaction. Appropriately diluted cDNA was later used for qPCR. RP49 was used as an internal control. Sequence of the primers used for PCR are as follows: *Oskar*: (*Osk_Forward primer – 5’ CTGAAGAACGGTCACCTCCT 3’; Osk_Reverse primer - 5’ GCCATTTGATGCAGCGAAAC 3’). Rp49: (RP4S_Fwd - 5’GGTATCGACAACAGAGTGCG 3’; Rp4S_Rev - 5’ATCTCCTTGCGCTTCTTGGA 3’)*.

### RNA Immunoprecipitation

*Drosophila* ovaries (100 pairs) from *rab7::YFP* and *w^1118^* animals were resuspended and homogenized in 300 μl of cold lysis buffer along with 2 μl RNase inhibitor from a 40U/ul (NEB#M0314S) stock and incubated on ice for 30 min. The lysate was centrifuged at 12,0000 rpm for 15 min. 30 μL of 10X DNase buffer (final conc. 1X) and 6 μl of DNase-I (Sigma#4716728001) from 10U/µl stock was added to the supernatant and incubated at 37°C for 30 min. The reaction was stopped by adding EDTA to a final concentration of 20 mM to stop the DNase activity. Total protein estimation was carried out by BCA kit (Pierce#23225). 10% of the total protein was aliquoted out as protein input; the remaining was used for IP and RNA isolation. For this, 1-2 mg of total protein was incubated with 3-5 μg of GFP-MBP-nanobody or MBP protein (control IP) overnight at 4 °C with gentle rotation. Following this step, 60 µL of Amylose beads (NEB#E8021S) was added to all the tubes and incubated for another 4 hours at 4° C with gentle rotation. Next, the beads were washed once with IP wash buffer, once with high salt buffer (IP wash buffer with 400 mM NaCl) followed by another wash with the IP wash buffer. The beads were then resuspended in 30 µL IP wash buffer and divided into 2 microcentrifuge tubes each with 10 µl and 20 µl of the beads respectively. The 10 µl aliquot was used for western blot to assess the immunoprecipation. The remaining 20 µl was processed for RNA isolation. 10 µl of 2x Proteinase-K buffer and 2.5 µl of Proteinase-

K (final conc. 2.5 µg/µl,) were added to the 20 µl bead volume and incubated at 55°c for 1hr to digest all the proteins to release the protein-bound RNAs. RNA was extracted using Trizol and chloroform extraction. RNA was precipitated from the aqueous layer using 600 µl of isopropanol together with 1.5 µl of glycogen (stock 20 mg/ml). 1µg of the RNA was used for cDNA synthesis in a 20 µl reaction volume as per the manufacturer’s instructions. Gene specific PCR (*osk*, *nos* and *gapdh*) was carried out using 6 µL of the cDNA. Primer sequences are as follows, *The Primer sequences are as follows, oskar : (Osk Forward primer: 5’ CGACAACGTGACGGATTTCCT 3’; Osk Reverse primer: 5’ GGAGGTGACCGTTCTTCAGG 3’), nanos: (Nos1F: 5’ CTCGTCGGCCACTTTGAGTC 3’ ; Nos1R: 5’ CTGTCGGCCAGAAAAGGGAAG 3’); gapdh (GAPDH_Fwd 5’ AAATCAAGGCTAAGGTCGAG 3’; GAPDH_Rev 5’CGAGATTAGCTTGACGAACT 3’)*.

## Contributions

AR conceptualized the project. VD, VK, SH, JS, TA and AR performed the experiments. VD, VK, SH, VS, JS, TA, GSR, VS and AR analyzed and interpreted the data. JS is deceased. JS gave consent to his authorship beforehand. AR wrote the initial draft; AR & GSR refined the subsequent drafts with inputs from all authors. AR led the project and acquired funding.

## Data G Resource availability

Raw data related to the images presented in the manuscript will be uploaded to a publicly shared server, in accordance with the journal’s guidelines.

## Acknowledgements

We thank the Daniel St. Johnston, Akira Nakamura, and Jo McDonald for generously sharing antibodies and fly lines; Mohit Kumar for fly lines, helpful discussions and troubleshooting. We thank the Bloomington *Drosophila* Stock Center (NIH P40OD018537) and Developmental Studies Hybridoma Bank, Iowa for fly stocks and antibodies. ARI confocal imaging facility and lab members for support.

## Supplementary Figures

**Supplementary Figure. 1.**
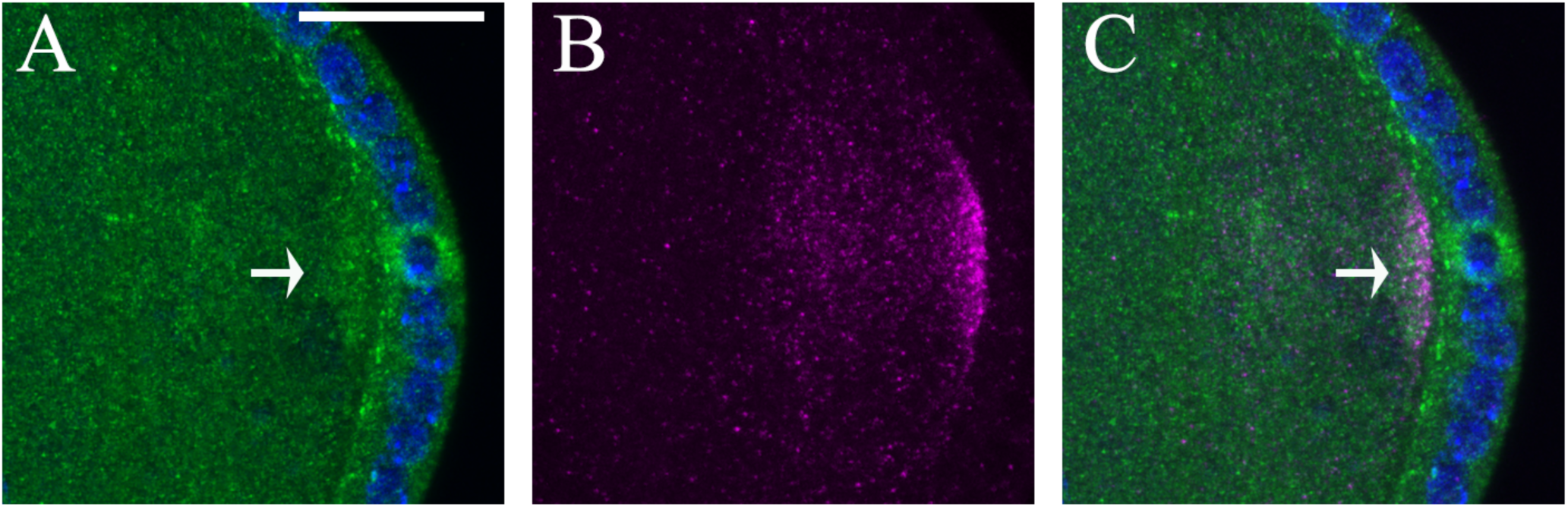
Localization of Stau with reference to Mon1:sfGFP. Stage 8 Mon1:sfGFP oocyte stained with anti-GFP (green), Staufen (magenta) and DAPI (blue). **(A)** Shown is the ‘platform-like’ Mon1:sfGFP staining seen near the posterior cortex. (B) Localization of Staufen. Not all Staufen is posteriorly localized at this stage. (C) Staufen staining is sandwiched between this ‘platform’ and the posterior cortex. Scale bar is 20μm.

**Supplementary Figure. 2.**
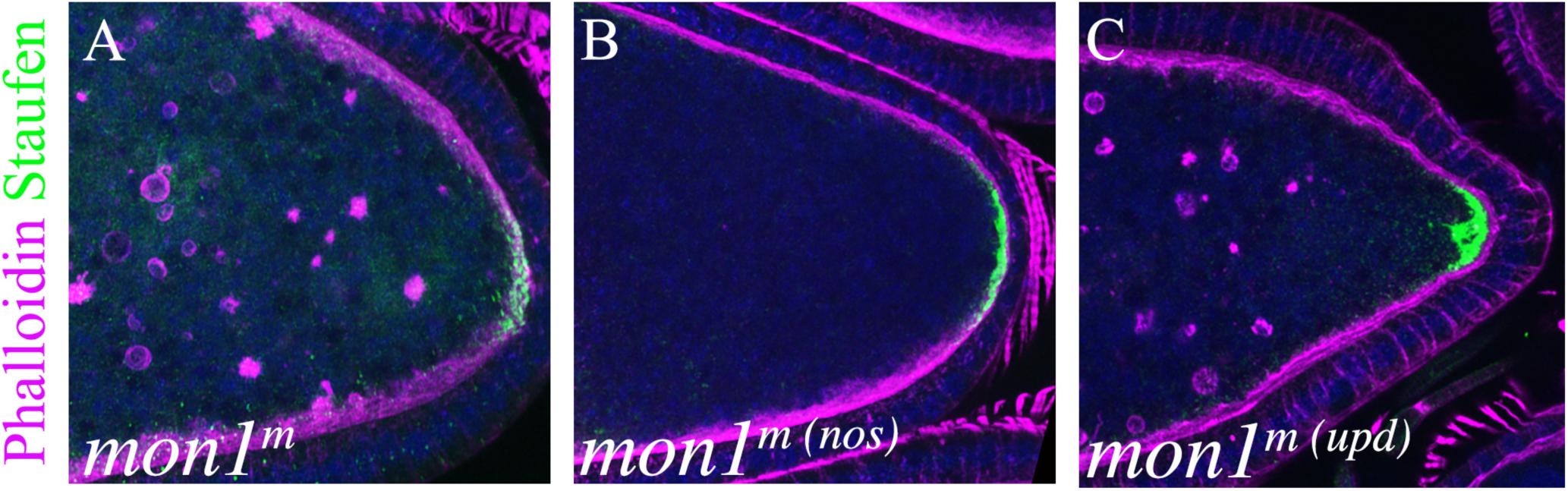
Actin rings in *mon1^m^* oocytes. Egg chambers stained with Staufen and Phalloidin **(A)** *mon1^m^ oocyte.* Actin rings (arrow) are seen in the ooplasm and budding from the lateral cortical membrane (asterisk). **(B)** *mon1^m (nos rescue)^*. Note absence of actin rings. **(C)** Actin rings persist in *mon^1m^* (upd rescue) oocytes.

**Movie S1**. Movie of showing mobility of control, *mon1^m^* and *mon1^m(rescue)^* larvae. Genotypes of the animals from left to right are as follows*: w^1118^*, *mon1^m^*, *mon1^m(nos rescue)^, mon1 ^(upd rescue)^ and mon1^m(c30C rescue)^*. *mon1^m(nos rescue)^* show normal crawling behaviour, comparable to wildtype.

## References

1. Assa-Kunik, E., I. L. Torres, E. D. Schejter, D. S. Johnston and B. Z. Shilo, 2007 Drosophila follicle cells are patterned by multiple levels of Notch signaling and antagonism between the Notch and JAK/STAT pathways. Development 134: 1161–1169.

2. Basargekar, A., S. Yogi, Z. Mushtaq, S. Deivasigamani, V. Kumar et al., 2020 Drosophila Mon1 and Rab7 interact to regulate glutamate receptor levels at the neuromuscular junction. Int J Dev Biol 64: 289–297.

3. Basu, U., S. S. Balakrishnan, V. Janardan and P. Raghu, 2020 A PI4KIIIalpha protein complex is required for cell viability during Drosophila wing development. Dev Biol 462: 208–222.

4. Bose, M., M. Lampe, J. Mahamid and A. Ephrussi, 2022 Liquid-to-solid phase transition of oskar ribonucleoprotein granules is essential for their function in Drosophila embryonic development. Cell 185: 1308–1324 e1323.

5. Bose, M., B. Rankovic, J. Mahamid and A. Ephrussi, 2024 An architectural role of specific RNA-RNA interactions in oskar granules. Nat Cell Biol 26: 1934–1942.

6. Brendza, R. P., L. R. Serbus, J. B. Duffy and W. M. Saxton, 2000 A function for kinesin I in the posterior transport of oskar mRNA and Staufen protein. Science 289: 2120–2122.

7. Cha, B. J., L. R. Serbus, B. S. Koppetsch and W. E. Theurkauf, 2002 Kinesin I-dependent cortical exclusion restricts pole plasm to the oocyte posterior. Nat Cell Biol 4: 592–598.

8. Cioni, J. M., J. Q. Lin, A. V. Holtermann, M. Koppers, M. A. H. Jakobs et al., 2019 Late Endosomes Act as mRNA Translation Platforms and Sustain Mitochondria in Axons. Cell 176: 56–72 e15.

9. Compagnon, J., L. Gervais, M. S. Roman, S. Chamot-Boeuf and A. Guichet, 2009 Interplay between Rab5 and PtdIns(4,5)P2 controls early endocytosis in the Drosophila germline. J Cell Sci 122: 25–35.

10. Deivasigamani, S., A. Basargekar, K. Shweta, P. Sonavane, G. S. Ratnaparkhi et al., 2015 A Presynaptic Regulatory System Acts Transsynaptically via Mon1 to Regulate Glutamate Receptor Levels in Drosophila. Genetics 201: 651-+.

11. Denef, N., and T. Schupbach, 2003 Patterning: JAK-STAT signalling in the Drosophila follicular epithelium. Curr Biol 13: R388–390.

12. Dhiman, N., K. Shweta, S. Tendulkar, G. Deshpande, G. S. Ratnaparkhi et al., 2019 Drosophila Mon1 constitutes a novel node in the brain-gonad axis that is essential for female germline maturation. Development 146.

13. Doerflinger, H., R. Benton, I. L. Torres, M. F. Zwart and D. St Johnston, 2006 Drosophila anterior-posterior polarity requires actin-dependent PAR-1 recruitment to the oocyte posterior. Curr Biol 16: 1090–1095.

14. Doerflinger, H., N. Vogt, I. L. Torres, V. Mirouse, I. Koch et al., 2010 Bazooka is required for polarisation of the Drosophila anterior-posterior axis. Development 137: 1765–1773.

15. Doerflinger, H., V. Zimyanin and D. St Johnston, 2022 The Drosophila anterior-posterior axis is polarized by asymmetric myosin activation. Curr Biol 32: 374–385 e374.

16. Dollar, G., E. Struckhoff, J. Michaud and R. S. Cohen, 2002 Rab11 polarization of the Drosophila oocyte: a novel link between membrane trafficking, microtubule organization, and oskar mRNA localization and translation. Development 129: 517–526.

17. Driever, W., and C. Nusslein-Volhard, 1988 A gradient of bicoid protein in Drosophila embryos. Cell 54: 83–93.

18. Ephrussi, A., and R. Lehmann, 1992 Induction of germ cell formation by oskar. Nature 358: 387–392.

19. Ferrandon, D., L. Elphick, C. Nusslein-Volhard and D. St Johnston, 1994 Staufen protein associates with the 3’UTR of bicoid mRNA to form particles that move in a microtubule-dependent manner. Cell 79: 1221–1232.

20. Gonzalez-Reyes, A., and D. St Johnston, 1994 Role of oocyte position in establishment of anterior-posterior polarity in Drosophila. Science 266: 639–642.

21. Jung, H., B. C. Yoon and C. E. Holt, 2012 Axonal mRNA localization and local protein synthesis in nervous system assembly, maintenance and repair. Nat Rev Neurosci 13: 308–324.

22. Kim-Ha, J., J. L. Smith and P. M. Macdonald, 1991 oskar mRNA is localized to the posterior pole of the Drosophila oocyte. Cell 66: 23–35.

23. Liao, Y. C., M. S. Fernandopulle, G. Wang, H. Choi, L. Hao et al., 2019 RNA Granules Hitchhike on Lysosomes for Long-Distance Transport, Using Annexin A11 as a Molecular Tether. Cell 179: 147-lpe120.

24. Logan, M. R., and C. A. Mandato, 2006 Regulation of the actin cytoskeleton by PIP2 in cytokinesis. Biol Cell 98: 377–388.

25. Long, R. M., R. H. Singer, X. Meng, I. Gonzalez, K. Nasmyth et al., 1997 Mating type switching in yeast controlled by asymmetric localization of ASH1 mRNA. Science 277: 383–387.

26. Lu, W., M. Lakonishok, R. Liu, N. Billington, A. Rich et al., 2020 Competition between kinesin-1 and myosin-V defines Drosophila posterior determination. Elife 9.

27. Mallart, C., S. Netter, F. Chalvet, S. Claret, A. Guichet et al., 2024 JAK-STAT-dependent contact between follicle cells and the oocyte controls Drosophila anterior-posterior polarity and germline development. Nat Commun 15: 1627.

28. Markussen, F. H., A. M. Michon, W. Breitwieser and A. Ephrussi, 1995 Translational control of oskar generates short OSK, the isoform that induces pole plasma assembly. Development 121: 3723–3732.

29. Mhlanga, M. M., D. P. Bratu, A. Genovesio, A. Rybarska, N. Chenouard et al., 2009 In vivo colocalisation of oskar mRNA and trans-acting proteins revealed by quantitative imaging of the Drosophila oocyte. PLoS One 4: e6241.

30. Micklem, D. R., J. Adams, S. Grunert and D. St Johnston, 2000 Distinct roles of two conserved Staufen domains in oskar mRNA localization and translation. EMBO J 19: 1366–1377.

31. Milas, A., J. de-Carvalho and I. A. Telley, 2023 Follicle cell contact maintains main body axis polarity in the Drosophila melanogaster oocyte. J Cell Biol 222.

32. Morais-de-Sa, E., A. Vega-Rioja, V. Trovisco and D. St Johnston, 2013 Oskar is targeted for degradation by the sequential action of Par-1, GSK-3, and the SCF(-)Slimb ubiquitin ligase. Dev Cell 26: 303–314.

33. Muntjes, K., S. K. Devan, A. S. Reichert and M. Feldbrugge, 2021 Linking transport and translation of mRNAs with endosomes and mitochondria. EMBO Rep 22: e52445.

34. Ramos, A., S. Grunert, J. Adams, D. R. Micklem, M. R. Proctor et al., 2000 RNA recognition by a Staufen double-stranded RNA-binding domain. EMBO J 19: 997–1009.

35. Ratnaparkhi, A., and K. Zinn, 2007 The secreted cell signal Folded Gastrulation regulates glial morphogenesis and axon guidance in Drosophila. Dev Biol 308: 158–168.

36. Riechmann, V., and A. Ephrussi, 2004 Par-1 regulates bicoid mRNA localisation by phosphorylating Exuperantia. Development 131: 5897–5907.

37. Riechmann, V., G. J. Gutierrez, P. Filardo, A. R. Nebreda and A. Ephrussi, 2002 Par-1 regulates stability of the posterior determinant Oskar by phosphorylation. Nat Cell Biol 4: 337–342.

38. Schuhmacher, J. S., S. Tom Dieck, S. Christoforidis, C. Landerer, J. Davila Gallesio et al., 2023 The Rab5 effector FERRY links early endosomes with mRNA localization. Mol Cell 83: 1839–1855 e1813.

39. Senju, Y., and P. Lappalainen, 2019 Regulation of actin dynamics by PI(4,5)P(2) in cell migration and endocytosis. Curr Opin Cell Biol 56: 7–13.

40. Shulman, J. M., R. Benton and D. St Johnston, 2000 The Drosophila homolog of C. elegans PAR-1 organizes the oocyte cytoskeleton and directs oskar mRNA localization to the posterior pole. Cell 101: 377–388.

41. Singer-Kruger, B., and R. P. Jansen, 2014 Here, there, everywhere. mRNA localization in budding yeast. RNA Biol 11: 1031–1039.

42. St Johnston, D., D. Beuchle and C. Nusslein-Volhard, 1991 Staufen, a gene required to localize maternal RNAs in the Drosophila egg. Cell 66: 51–63.

43. St Johnston, D., and C. Nusslein-Volhard, 1992 The origin of pattern and polarity in the Drosophila embryo. Cell 68: 201–219.

44. Tanaka, T., Y. Kato, K. Matsuda, K. Hanyu-Nakamura and A. Nakamura, 2011 Drosophila Mon2 couples Oskar-induced endocytosis with actin remodeling for cortical anchorage of the germ plasm. Development 138: 2523–2532.

45. Tanaka, T., and A. Nakamura, 2008 The endocytic pathway acts downstream of Oskar in Drosophila germ plasm assembly. Development 135: 1107–1117.

46. Tanaka, T., and A. Nakamura, 2011 Oskar-induced endocytic activation and actin remodeling for anchorage of the Drosophila germ plasm. Bioarchitecture 1: 122–126.

47. Tanaka, T., N. Tani and A. Nakamura, 2021 Receptor-mediated yolk uptake is required for oskar mRNA localization and cortical anchorage of germ plasm components in the Drosophila oocyte. PLoS Biol 19: e3001183.

48. Tomancak, P., F. Piano, V. Riechmann, K. C. Gunsalus, K. J. Kemphues et al., 2000 A Drosophila melanogaster homologue of Caenorhabditis elegans par-1 acts at an early step in embryonic-axis formation. Nat Cell Biol 2: 458–460.

49. Vanzo, N., A. Oprins, D. Xanthakis, A. Ephrussi and C. Rabouille, 2007 Stimulation of endocytosis and actin dynamics by Oskar polarizes the Drosophila oocyte. Dev Cell 12: 543–555.

50. Vanzo, N. F., and A. Ephrussi, 2002 Oskar anchoring restricts pole plasm formation to the posterior of the Drosophila oocyte. Development 129: 3705–3714.

51. Yousefian, J., T. Troost, F. Grawe, T. Sasamura, M. Fortini et al., 2013 Dmon1 controls recruitment of Rab7 to maturing endosomes in Drosophila. J Cell Sci 126: 1583–1594.

52. Yu, Y., D. Chen, S. M. Farmer, S. Xu, B. Rios et al., 2024 Endolysosomal trafficking controls yolk granule biogenesis in vitellogenic Drosophila oocytes. PLoS Genet 20: e1011152.

